# SimpylCellCounter: An Automated Solution for Quantifying Cells in Brain Tissue

**DOI:** 10.1101/2020.02.22.960948

**Authors:** Aneesh Bal, Fidel Maureira, Amy A. Arguello

**Affiliations:** Psychology Dept, Michigan State University, Giltner Hall, Rm 213A, East Lansing, MI, 48825, USA; Biological Systems Engineering, Washington State University, Paccar 351 Pullman, WA, 99164-6120, USA

**Author notes:** <u>Corresponding Author</u>: Dr. Amy A. Arguello, Michigan State University, Department of Psychology, Behavioral Neuroscience, Giltner Hall, Rm 213A, 293 Farm Lane, East Lansing, MI 48825.

**Keywords:** Automated quantification, convolutional neural network, Cfos, brain tissue

## Abstract

**Rationale & Objective:** Manual quantification of activated cells can provide valuable information about stimuli-induced changes within brain regions; however, this analysis remains time intensive. Therefore, we created SimpylCellCounter (SCC), an automated method to quantify cells that express Cfos protein, an index of neuronal activity, in brain tissue and benchmarked it against two widely-used methods: OpenColonyFormingUnit (OCFU) and ImageJ Edge Detection Macro (IMJM).

**Methods:** In Experiment 1, manually-obtained counts were compared to those detected via OCFU, IMJM and SCC. The absolute error in counts (manual *versus* automated method) was calculated, and error types were categorized as false positives or negatives. In Experiment 2, performance analytics of OCFU, IMJM and SCC were compared. In Experiment 3, SCC performed analysis on images it was not trained on, to assess its general utility.

**Results & Conclusions:** We found SCC to be highly accurate and efficient in quantifying both cells with circular morphologies and those expressing Cfos. Additionally, SCC utilizes a new approach for counting overlapping cells with a pretrained convolutional neural network classifier. The current study demonstrates that SCC is a novel, automated tool to quantify cells in brain tissue, complementing current, open-sourced quantification methods designed to detect cells *in vitro*.

## INTRODUCTION

Immediate early genes (IEGs) are rapidly transcribed and translated upon stimulus exposure, making them useful for post-behavioral, correlated readouts of cellular activity^1^. Cfos, a commonly studied IEG, is a proto-oncogene and member of the Fos family of transcription factors. *cfos* mRNA is transcribed within minutes of stimulus exposure and results in the cytoplasmic expression of Cfos protein 60-90 minutes later^2^. Behaviorally-induced increases in the number of Cfos-Immunoreactive (Cfos-IR) cells often suggest that these activated neuronal ensembles may contribute to specific behaviors. For example, characterization of the brain regions and cell types in which Cfos protein is increased has provided insight into the cell populations that contribute to learning and memory, drug addiction, obesity and fear conditioning^3–11^.

To analyze Cfos-IR cells, experimenters can choose between manual or automated methods for quantification. Brain slices are typically immunohistochemically stained for Cfos protein, resulting in round, light to dark-labeled cells. For manual quantification, cells can be counted in areas within a set microscopic field of view, or digital images of Cfos-stained tissue are obtained, imported and thresholded using software such as ImageJ to aid in counting^4,5,12,13^. While analysis with select software improves the reliability of manual quantification, an experimenter must still determine whether to count a dark-stained cell based on smoothness, clarity and size. Additionally, as the number of images increases, so does fatigue and the potential for increased errors in counts. Alternatively, existing automated methods can be used to quantify Cfos-like cells even though they are optimized for cell colony or tumor spheroid analysis. However, these algorithms can encounter problems with edge detection, contrast enhancement and denoising in brain tissue analysis^14–18^. Edge detection allows for clean segmentation of cells within colonies; however, this algorithm can detect false edges in background staining present with Cfos cells, which can obscure the cells of interest. While contrast enhancement makes it easier to detect Cfos-IR cells, it can result in an overestimation of total cell count due to increased pixel intensity of dimly-stained cells. Lastly, denoising can remove background noise from Cfos images, but it can also lead to false negatives in images where Cfos-IR cells may be slightly out-of-focus. Therefore, to increase objectivity, reliability and minimize the time required to analyze images, an improved automated method for quantifying Cfos-like cells in brain tissue is required.

We created SimpylCellCounter (SCC), an efficient and accurate automated method for quantifying Cfos-IR cells in brain tissue. SCC utilizes binary thresholding and morphological functions from the open-sourced computer vision library OpenComputerVision^19^, implemented in Python. SCC allows a user to manually set parameters of darkness with basic thresholding, cell size and circularity by filtering out non-circular objects and counting only user-defined objects (Fig 1). SCC also utilizes OpenCV’s highly-efficient, image processing functions to rapidly batch process large sets of digital images and incorporates a new approach to separating overlapping cells via a convolutional neuronal network.

**Figure 1:**
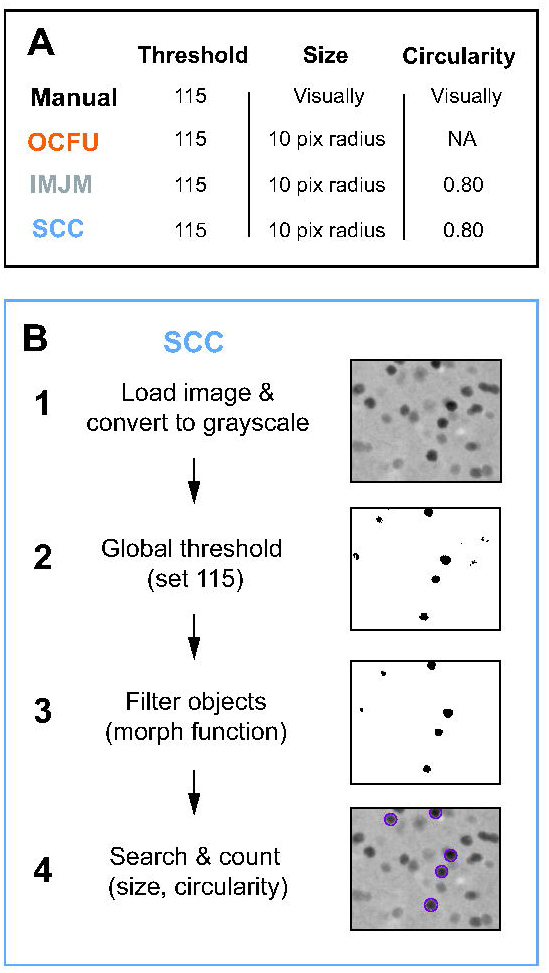
Schematic of Automated Processing Steps. **A)** Selected parameters for threshold (pixel intensity), object size (pixel radius) and object circularity for Manual, OpenColonyFormingUnit (OCFU), ImageJ Edge Detection Macro (IMJM) and SimpylCellCounter (SCC) of Cfos-immunoreactive (Cfos-IR) cells. **B)** Image processing sequence for SCC: 1) loads images and converts to 8-bit grayscale making all pixel intensities range between 0 and 255, 2) performs global threshold based on a user-set value and creates a binarized mask, 3) performs morphological operations on the binary mask to filter out noise-like particles, 4) further selects for objects based on size, circularity and cell overlap, leading to a final cell count.

To test the feasibility and efficiency of SCC, we compared our algorithm to two, highly-cited, open-sourced, cell colony-based automated quantification methods: OpenColonyFormingUnit^17^ (OCFU) and ImageJ Edge Detection Macro^16^ (IMJM). We chose OCFU and IMJM due to the similarities between Cfos-IR cells and the cell colony images analyzed in their respective publications. We used 192 images of Cfos-IR cells from the orbitofrontal cortex (OFC) of rats that underwent a cue-induced reinstatement paradigm where previously drug-paired cues elicited increased drug-seeking behaviors. We tested various metrics of performance between OCFU, IMJM and SCC and found that SCC quantified Cfos-IR cells with high accuracy when compared with manual analysis. SCC displayed the fastest quantification time of all automated methods tested and maintained accuracy and efficiency when threshold values and image size were changed. Lastly, we showed that SCC generalized across multiple sets of images (fabricated Cfos images, *S. aureus* and *E. coli* colonies), indicating that it was not overfit to our laboratory’s method of Cfos analysis.

## RESULTS

### Comparison of Cfos-IR counts between manual and automated methods (EXP 1)

The number of Cfos-IR cells at several bregma points within the ventral OFC (vOFC) was quantified manually (white) or with three automated methods: OCFU (orange), IMJM (gray) or SCC (blue) (Fig 2). Specifically, a 4×6 ANOVA revealed no significant Method x Bregma interaction effect or main effects of Method or Bregma (Fig 2A). Therefore, all automated methods (OCFU, IMJM and SCC) displayed similar average number of Cfos-IR cells at each bregma point. An ANOVA of the total number of vOFC Cfos-IR cells quantified by each method did not reveal a significant effect. However, there was a trend (F_3,28_ = 2.51, p = 0.079) for an increased total number of cell counts by OCFU and IMJM, but not SCC, compared to manual. (Fig 2B).

**Figure 2:**
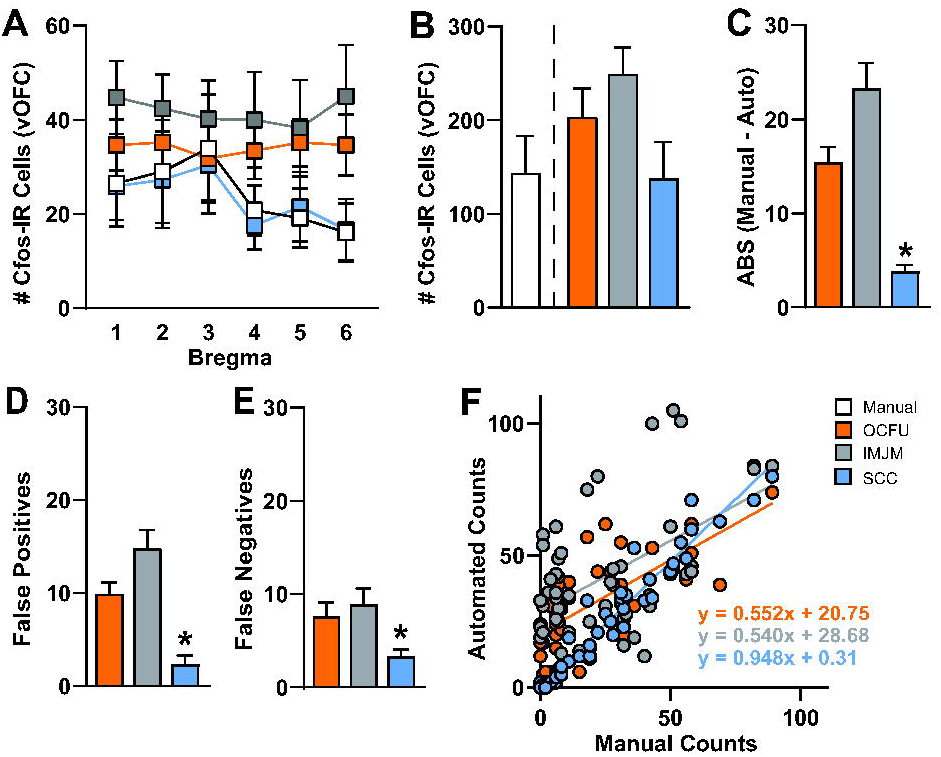
Comparison of Manual *vs* Automated Quantification Methods. Manual *vs* automated quantification of Cfos-IR cells obtained from the ventral orbitofrontal cortex (vOFC) of rats that underwent cue-induced reinstatement of drug-seeking behavior. The automated methods included: OCFU (orange), IMJM (gray) and SCC (blue). **A)** Average number of Cfos-IR cells counted by manual *vs* automated methods over several points along the anterior-posterior axis (bregma + 5.12 to + 3.72) of the vOFC, n = 96 total images per method. **B)** Average number cell counts. **C)** Average absolute error: *ABS(Manual counts - Automated counts)*, **D)** Average number of false positives, number of cells detected by automated methods that were not counted manually, n = subset of 30, **E)** Average number of false negatives, number of cells counted manually that were not detected by automated methods, n = subset of 30 images, **F)** Correlation of manual *vs* automated counts. Manual correlated with OCFU, p < 0.001 and regression of y = 0.552x + 20.75; Manual correlated with IMJM, p < 0.001 and regression of y = 0.540x + 28.68; Manual correlated with SCC, p < 0.001 and regression of y = 0.948x + 0.31.

Next, we calculated the difference in the number of Cfos-IR cells counted between manual and each automated method. ANOVA of the total absolute errors per method revealed a significant effect (F_2, 141_ = 30.41, p<0.0001). Post-hoc analysis revealed that the absolute error between manual and automated counts determined by SCC was significantly less than both OCFU and IMJM (Bonferroni’s test, p<0.01). Therefore, compared to IMJM and OCFU, SCC had the least difference in cell counts compared to manual analysis (Fig 2C).

To further explore the types of errors, we conducted an analysis of false positives (Fig 2D) and false negatives (Fig 2E) for all automated methods. ANOVA of false positives revealed a significant effect, (F_2,87_ = 22.37, p<0.0001), with SCC displaying a significantly lower magnitude of false positives compared to OCFU and IMJM (Bonferroni’s test, p<0.01). Additionally, an ANOVA of false negatives revealed a significant effect, (F_2,87_ = 5.27, p<0.01) with SCC detecting significantly lower magnitude of false negatives compared to IMJM (Bonferroni’s test, p<0.01) but not OCFU. Therefore, SCC minimized detection of false positives and negatives, resulting in a smaller number of absolute errors compared to OCFU and IMJM. Examples of types of false positives (plus symbol) and negatives (carrot symbol) compared to manual counts (magenta circles) are depicted for each automated method (Fig 3).

**Figure 3:**
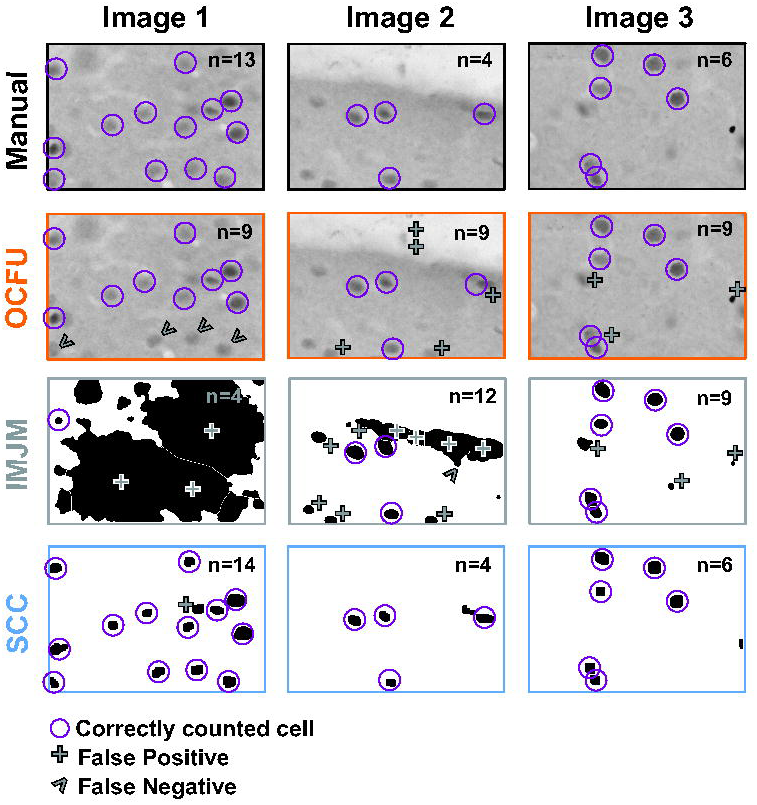
Characterization of Automated Quantification. Comparison of 3 images with manual *vs* automated method with examples of count classification: OCFU (orange), IMJM (gray) and SCC (blue). The number of detected cells is displayed in the upper right corner of each image. Correctly counted cells (compared to manual) are depicted in magenta circles. False positives are depicted by a plus symbol while false negatives are depicted by the carrot symbol.

Lastly, the number of Cfos-IR cells quantified with SCC was correlated with manual counts (Fig 2F). Linear regression analysis of manual *vs* automated counts revealed the following: manual *vs* OCFU, p<0.0001 with a regression equation of y = 0.552x + 20.75; manual *vs* IMJM, p<0.0001 with a regression equation of y = 0.540x + 28.68; manual *vs* SCC, p<0.0001 with a regression equation of y = 0.948x + 0.31. Therefore, SCC detected similar cell counts per image when compared to manual analysis.

### Differences in automated method performance (EXP 2)

We compared the performance analytics of OCFU, IMJM and SCC at analyzing Cfos-IR cells (Fig 4). We determined the average time (sec) for each automated method to quantify one image (Fig 4A). An ANOVA of time per method revealed a significant effect (F_2,87_ = 1292, p<0.0001), with SCC exhibiting the lowest analysis time (Bonferroni’s test, p<0.01). The time required to quantify a set of images as a function of image size was also compared (Fig 4B). A 3×7 repeated measures ANOVA of analysis time per method revealed a significant Method x Size interaction effect (F_12,522_ = 2271, p<0.0001) and a significant main effect of Size (F_6,522_=9773, p<0.0001), with SCC exhibiting the lowest time to analyze one image across size groups (Bonferroni’s test, p<0.01). Therefore, all 3 automated methods display increases in processing time with larger image size. While SCC’s processing speed increases as a function of image size, it exhibits the most rapid analysis compared to OCFU and IMJM.

**Figure 4:**
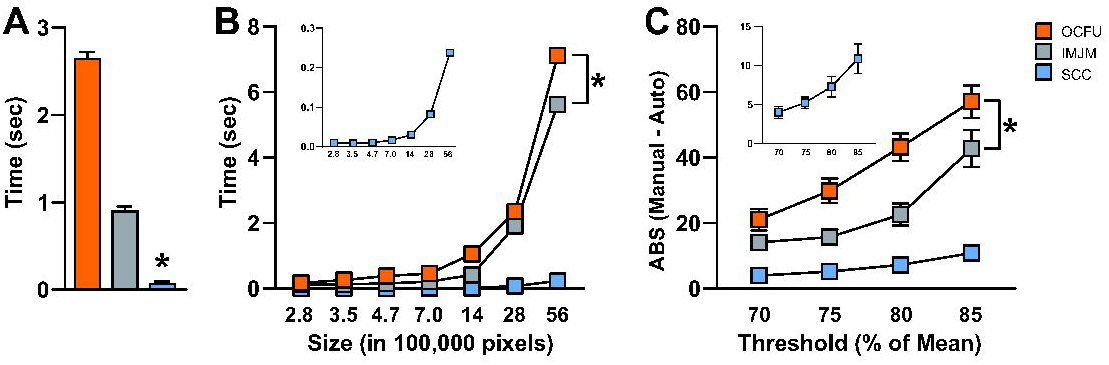
Performance Analytics of Automated Methods. OCFU (orange), IMJM (gray) and SCC (blue) performance analytics were compared. **A)** Average time (sec) to quantify Cfos-IR cells per image (1920 x 1460 pixels), n = 30 images per automated method. **B)** Average time (sec) as a function of image size (pixels) to quantify the number of Cfos-IR cells per image, n = 30 images per automated method. Inset displays data for SCC only. **C)** Average absolute error per image where: *ABS(Manual counts - Automated counts)* as the binary threshold value approaches the mean pixel intensity of the image, n = 15 images per threshold factor. Inset displays data for SCC only.

Lastly, we compared the absolute error of each automated method as the threshold varied as a percentage of mean pixel intensity per image (Fig 4C). A 3×4 repeated measures ANOVA of absolute errors per method revealed a significant Method x Threshold group interaction effect (F_6,126_ = 9.11, p<0.0001) and a significant main effect of Threshold group (F_3,126_ = 66.08, p<0.0001), with SCC displaying the lowest absolute error across threshold groups (Bonferroni’s test, p<0.01). Therefore, SCC displays robust accuracy even as threshold percentage and background noise increases.

### Determining SCC’s performance on different image types (EXP 3)

We compared ground truth (white) to SCC (blue) counts on three different types of images: fabricated Cfos-images, *S. aureus* and *E. coli* cell colonies (Fig 5). Independent samples t-test of average ground truth counts *vs* SCC counts revealed no significant differences (Fig 5B, t_28_ = 0.13, p = 0.89). Linear regression of ground truth *vs* SCC counts revealed a significant correlation (Fig 5C, p<0.0001) and regression equation of y = 0.991x – 0.82. Independent samples t-test of average ground truth counts *vs* SCC counts revealed no significant differences between groups (Fig 5E, t_26_ = 0.11, p = 0.91). Linear regression of ground truth *vs* SCC counts revealed a significant correlation (Fig 5F, p<0.0001) and regression equation of y = 0.968x + 0.10. Independent samples t-test of average ground truth counts *vs* SCC counts revealed no significant differences between groups (Fig 5H, t_28_ = 0.20, p = 0.84). Linear regression of ground truth *E. coli* counts *vs* SCC counts revealed a significant correlation (Fig 5I, p<0.0001) and regression equation of y = 0.943x + 0.39. Taken together, these results show that SCC counts matched ground truth counts for fabricated Cfos images (Fig 5A-C), *S. aureus* images (Fig 5D-F) and *E. coli* colonies (Fig 5G-I). Therefore, SCC accurately quantifies cell types that it was not trained on, indicating generalizability.

**Figure 5:**
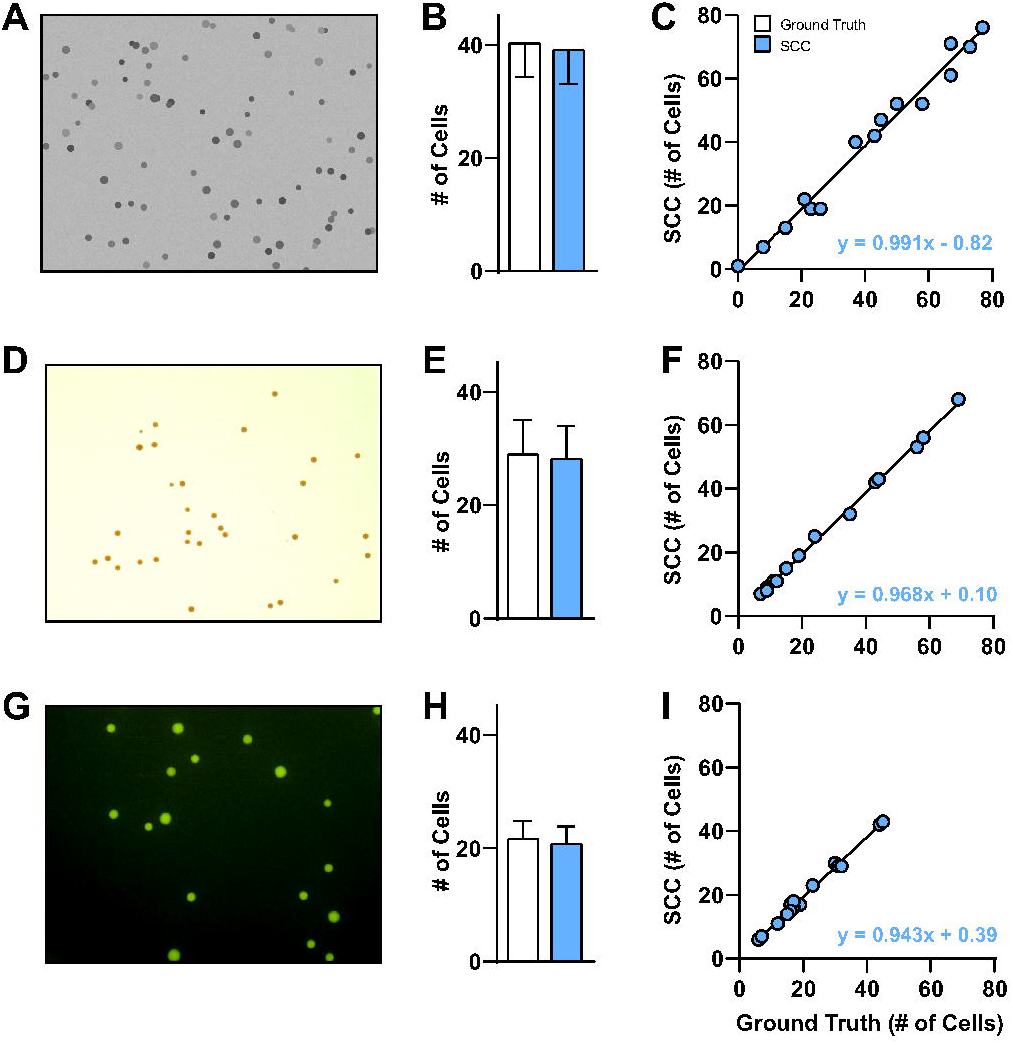
Overfitting Analysis. SCC evaluated multiple sets of data including fabricated Cfos images and non-neuronal cell types *S. aureus* and *E. coli* to test the generalizability of our algorithm. Ground truth was defined as: the number of known Cfos-IR cells that meet threshold, size, and circularity criterion on fabricated Cfos images (**A-C**) and the number of cells per image counted by OCFU (**D-I**). OCFU was trained on these two datasets, meaning it is accurate, thereby making it “ground truth”. Ground truth (white bars), SCC (blue bars). **A)** Representative image of fabricated Cfos cells, **B)** Average number of cells counted by ground truth *vs* SCC, n = 15 images, **C)** Ground truth counts correlated with SCC counts resulted in p < 0.001 and regression of y = 0.991x - 0.82, **D)** Representative image of *S. aureus*, **E)** Average number of cells counted by manual *vs* SCC, n = 14 images, **F)** Ground truth counts correlated with SCC counts resulted in p < 0.001 and regression of y = 0.968x + 0.10, **G)** Representative image of *E. coli*, **H)** Average number of ground truth cells *vs* SCC counts, n = 15 images, **I)** Ground truth counts correlated with SCC counts resulted in, p < 0.001 and regression of y = 0.943x + 0.39.

## DISCUSSION

The present study aimed to create an automated method to analyze cell number in brain tissue, to complement existing open-sourced methods designed for cell colony analysis. We created SimpylCellCounter (SCC) and used this automated method to quantify the number of Cfos-immunoreactive (IR) cells in brain tissue. We analyzed several variables and found that SCC 1) detected a similar magnitude of cells as manual analysis, 2) displayed low absolute errors, false positives and negatives compared to two widely-used automated methods, OpenColonyFormingUnit (OCFU) and ImageJ Edge Detection Macro (IMJM) and 3) was rapid at processing images of increasing size and 4) detected similar number of cells across varying thresholds suggesting that this algorithm can maintain accuracy when this parameter changes. Importantly, SCC introduces a novel approach to detect and quantify overlapping cells with the use of Hu Moments and a convolutional neural network (CNN). Use of a pretrained CNN affords rapid analysis along with high levels of accuracy at quantifying overlapping cells. For binarized objects with significant overlap, it is difficult to judge whether there are multiple cells or simply one cell with irregular morphological features. Our CNN implementation is optimal for such a task since it learns morphological features of binarized single or multiple cells, making input data classification with a pretrained network a quick process.

SCC-driven analysis of Cfos-IR cell number was correlated with manual counts. We previously found that rats presented with a drug-associated cue displayed an increased number of Cfos-IR cells in the ventral orbitofrontal cortex (vOFC). Specifically, when compared to previous manually-analyzed data, SCC detected similar numbers of total Cfos-IR cells (Fig 2B) and similar numbers of counts in anterior-posterior divisions of the OFC (Fig 2A). Furthermore, there is a near perfect match of cell counts analyzed manually *vs* with SCC (Fig 2F). Taken together, these data indicate that SCC is comparable to manual analysis, accurate at providing objective estimates of cell number and efficient since less time is needed to determine whether a cell should be counted based on size, shape and pixel intensity.

We also compared manual analysis of cell counts to two, commonly-used algorithms OCFU and IMJM, which analyze cell number in colonies but can also be used for other applications requiring detection of circular objects. We did not benchmark SCC against general cell analysis methods like CellProfiler^20^ and Ilastik^21^, since they require the user to have a working understanding of computer vision algorithms to create a custom pipeline for analysis prior to inputting data, making these solutions less user-friendly.

We wanted to examine whether OCFU, IMJM and SCC detected similar numbers of cells. While not significant, there was a trend for OCFU and IMJM to result in higher total numbers of Cfos-IR cells in the vOFC and across bregmas, compared to manual and SCC (Fig 2B). When we compared the absolute error between manual and automated counts, we found that SCC resulted in significantly less errors than OCFU and IMJM (Fig 2C). We then aimed to understand the potential reasons for differences in cell number between manual, OCFU, IMJM and SCC methods. Examples of false negatives occurred where OCFU filtered out Cfos-IR cells that were oblong shaped or blurry. False positives may have occurred when cells displayed a color gradient (half of cell dark, other half light), resulting in a cell being counted twice (Fig 3, **OCFU**). Examples of false negatives in IMJM occurred when contrast enhancement created a loss of difference in pixel intensity in a cell *vs* background, resulting in edge detection failure and omission of neighboring positive cells. Additionally, filtering procedures may alter cell morphology, making it difficult for IMJM’s watershed algorithm to effectively separate certain overlapping cells, leading to lower counts. (Fig 3, **IMJM**). False positives could result from errors where background, noise-like particles with correct cell shape are erroneously filled with IMJM’s “fill holes” step, leading to increased counts.

SCC is a brain-specific algorithm that complements currently-available, automated quantification methods, offering improvements in speed of digital analysis. SCC’s simple processing scheme only includes functions that are essential to separating Cfos-IR cells from background and noise, such as thresholding, dilation and erosion. Similar to OCFU and IMJM, SCC initially processes an entire digital image, until step 4 of the algorithm (Fig 1A), when it then computes Hu Moments to independently and sequentially quantify circularity in a contour-wise fashion. Therefore, SCC decides which objects demand additional time for analysis: non-circular contours are input into the CNN to test for overlapping cells while circular objects are simply counted, thereby reducing processing times (Fig 4A). Furthermore, we also determined that SCC is consistently faster at processing increasingly larger images (Fig 4B) and maintains a high level of count accuracy even as threshold values change (Fig 4C). As thresholds approach the mean pixel intensity of the image being processed, the absolute error (manual – automated) consistently increases for all automated methods. However, SCC is able to minimize this error, likely due to the dynamic filtering operations: as threshold approaches the mean, objects are more rigorously filtered by increasing iterations of dilation and erosion.

Lastly, we aimed to determine whether the SCC algorithm could also accurately analyze cell counts from other lab data. To do this, we utilized data from artificially-constructed Cfos-IR images that contained cells in varying size and intensity. SCC exhibits accurate performance, as shown by raw averages and correlation (Fig 5A-C). Furthermore, we utilized sample images obtained from the OCFU database from *S. aureus* and *E. coli* image samples. SCC displayed strong correlations between manual *vs* automated detection method, demonstrating that our algorithm can detect circular, non-Cfos-IR, objects including cells that are pigmented and from *in vitro* mediums (Fig 5D-F, G-I). These data demonstrate that SCC is not overfit to the data it was trained on and can likely generalize to other datasets collected by different labs. With that being said, even though SCC can count other types of cells in bacterial cultures, SCC is built for Cfos-like images.

In the future, SCC can potentially be used to effectively analyze viral or fluorescent images. For example, colorimetric Cfos images have dark cells and a lighter background, whereas fluorescently-labeled cells would be lighter than the background. Therefore, the user can simply invert the threshold in SCC and proceed with the same processing chain. Given that the SCC code is flexible, a user can easily adjust parameters of threshold, size and filtering to fit their specific application. SCC is an accurate, efficient and novel automated tool to quantify Cfos-like cells in brain tissue and can be extended to analysis of cells with circular morphologies.

## MATERIALS & METHODS

### Animals and Behavioral Experiments

We utilized a previously published data set in which the number of Cfos-IR cells was increased with exposure to cocaine-associated cues^22^. Male Sprague-Dawley rats (Envigo Inc, Haslett, MI, N=20) were housed under reversed lighting conditions (lights off 7am, on 7pm) and were fed 20-25g of standard irradiated rodent chow with water available *ad libitum*. Protocols were approved by the Institutional Animal Care and Use Committee (IACUC) at Michigan State University (MSU) and followed the National Research Council’s Guide for the Care and Use of Laboratory Rats. Intravenous catheters were subcutaneously implanted into the right jugular vein. Following 5 days of recovery, rats underwent cocaine self-administration and extinction training, followed by drug-seeking tests without cue (EXT) or with cue presentation (TEST). After tests, rats were sacrificed and perfused, brains were extracted and cryoprotected and sectioning and image processing was conducted as previously described^22^. We obtained a total of 192 images of Cfos-stained sections from lateral and ventral OFC (lOFC, vOFC respectively) from six bregma points spanning the A-P axis (+5.12 to +3.72), from which Cfos-IR cells were quantified^22^. In the current study, we compared counts that were previously analyzed manually, with the automated methods OCFU, IMJM and SCC.

### Image Processing Steps and Parameter Selection

#### Manual

We imported images into ImageJ*v*1.51^13^, converted them to 8-bit grayscale and applied threshold values with the top value set to 115 and the bottom set to 120. These adjustments created a round, red contour around the darkest cells on each image, which assisted experimenters to judge the circularity and size of cells during counting (Fig 1A).

#### OCFU

We utilized the OCFU graphic user interface application for all experiments. We set two parameters: radius size (minimum at 10 pixels, maximum at auto-max) and threshold to 61. Since OCFU’s threshold does not directly correspond to pixel intensity, we incorporated a standardization procedure (Supplementary Methods & Supplementary **Fig S2**) by which a threshold value of 61 in OCFU was equivalent to the value of 115 in manual, IMJM and SCC. (Fig 1A).

#### IMJM

This algorithm contained numerous parameters that were local to ImageJ functions but not common to OCFU or SCC. Therefore, we used parameters for IMJM provided in^16^. We modified the contrast enhancement value to 0.001 and circularity value to 0.8-1.0 (Fig 1A). We also added a binary threshold step and implemented IMJM in MATLAB’s ImageJ wrapper, MIJI^23^. The exact MIJI workflow can be found at: https://github.com/aneeshbal/SimpylCellCounter/blob/master/recreationFunctions/AutoQMS_MIJI.m

#### SCC

##### Binary Mask

In the first processing step, the user selects the folder of images to be analyzed then SCC applies a binary threshold to 8-bit images, converting all pixel values lower than the set threshold to black and those higher than threshold to white. After thresholding, shapeless, poorly-connected, sparse groups of black pixels represent background, noise-like particles while round, well-connected, dense collections of black pixels represent Cfos-IR cells.

##### Dilation & Erosion

In the dilation step, white pixels (background) engulf adjacent black pixels (cells of interest + noise). Consequently, small, noise-like objects are completely engulfed by white pixels and become part of the background. To recover an object’s original morphology, which is altered with dilation, SCC performs an erosion step (opposite of dilation). Dilation and erosion steps occur for a set number of iterations, determined by the user-set threshold to mean pixel intensity ratio (MPI). As the threshold-MPI ratio approaches 1, the magnitude of noise-like particles exponentially increases, therefore SCC accordingly increases the iterations of these steps resulting in a stringent filtering process.

##### Object Selection

Following dilation and erosion, SCC discards objects based on size criteria by drawing contours over all the objects on the filtered binary mask, calculates the zeroth-order moment of each contour (area) and discards all contours with a smaller area than the user-set criteria (pixel radius converted to area). Following this step, certain objects may be overlapping, obscuring the total cell count. Rather than performing the popular watershed segmentation algorithm to separate overlapping objects, which can alter cell morphology^24^, we utilized Hu Moments to compute contour circularity^25,26^. Hu Moments are orientation- and scale-invariant properties intrinsic to shapes. Perfectly circular contours resulted in a log-adjusted first Hu Moment value of ~0.79. We observed that a single contour surrounds the perimeter of overlapping objects, resulting in a non-circular contour with a first Hu Moment value typically below ~0.76. SCC then applies a pretrained convolutional neural network (CNN) classifier (Supplementary Methods & Supplementary **Fig S1**) to determine the number of cells within non-circular contours and adds these to the number of circular contours. For all experiments, radius size was set to 10 pixels, threshold was 115 and circularity was 0.7 unless otherwise stated. (Fig 1A).

### Experiment 1: Accuracy and Feasibility of Automated Methods

Using 192 images of vOFC brain sections from TEST rats, we compared the average number of Cfos-IR cells per bregma point, and the total number of Cfos-IR cells. Next, we calculated the absolute error of cell counts between manual *vs* each automated method (OCFU, IMJM, SCC) and then conducted an error analysis in a subset of images. For each manually-counted cell, the number of false positives (cells counted by automated methods but not manually) and false negatives (cells counted manually but not by automated methods) was determined. Last, we correlated the number of counts detected via manual *vs* each automated method and conducted a linear regression analysis.

### Experiment 2: Performance Analytics of Automated Methods

Using 30 randomly selected Cfos-IR images from EXT and TEST subjects (lOFC and vOFC), we calculated the average time (sec) to process one image (1920 x 1460 pixels), then resized each image (by factors of 0.5, 1, 2, 4, 6, 8, 10) and calculated the resulting image size: 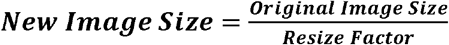. For each resize factor, we calculated the average time (sec) for each automated method to process 30 images. Lastly, using 15 randomly-selected Cfos-IR images from EXT and TEST subjects (lOFC and vOFC), we calculated the absolute error for each automated method as a function of a changing threshold. For each image, we multiplied the MPI by each threshold factor (0.7, 0.75, 0.8 or 0.85) to obtain the final threshold value. We then quantified Cfos-IR cells and the absolute error for each image between manual *vs* automated methods. We did not use the command line interface for OCFU but instead quantified the image processing time on the graphic user interface (GUI) from the input image until a cell count was displayed.

### Experiment 3: Overfitting Analysis for SCC

We obtained 3 separate datasets of non-Cfos images including: 1) fabricated Cfos images (n=15), 2) *S. aureus* colony images (n=14) and 3) *E. coli* colony images (n=15). We created 15 fabricated Cfos images, using a Python implementation of OpenCV by placing a random number of circles (between 5 and 100) of varying pixel intensities and sizes on a gray background that closely resembles the background staining of Cfos images. Additionally, we obtained cell colony images of *S. aureus* and *E. coli* from the open-sourced database provided by Dr. Quentin Geissman at the following link: http://opencfu.sourceforge.net/samples.php. The source images contain agar plates but since SCC does not have a region of interest selector, images were cropped to include only a subset of contents inside agar plates. The final images used are provided at: https://github.com/aneeshbal/SimpylCellCounter/tree/master/imageSamples.

For fabricated Cfos-IR images, we calculated ground truth counts and compared them to SCC counts. Ground truth here was defined as the number of cells that met the user-defined, size and threshold criteria, and since these images were fabricated, the exact number of cells was pre-determined. We calculated the average counts per image and performed a correlation and linear regression of ground truth *vs* SCC counts. For cell colony images, we defined ground truth as the number of cells counted by OCFU, given that it was optimized for cell colony images. We then repeated the analyses performed on the fabricated Cfos-IR images on *S. aureus* and *E. coli* image samples.

### Code Availability

All code is available: https://github.com/aneeshbal/SimpylCellCounter

### Statistics

For EXPs 1 & 2, repeated measures analysis of variance (ANOVA) were conducted for mean cell counts across bregma points comparing manual *vs* automated methods (within-subject factor = bregma, between-subject factor = method), time per automated method across image size groups (within-subject = image size, between-subject = method) and absolute errors by automated methods across threshold factor (within-subject = threshold group, between-subject = method). One-way ANOVAs were conducted to explore differences in total cell counts, absolute errors, false positives and negatives across automated methods and time taken to analyze images per automated method. Additionally, linear regression was conducted to examine correlations between manual *vs* automated counts and ground truth *vs* SCC counts. For EXP 3, independent t-tests were conducted to examine differences between ground truth *vs* SCC counts. For all statistical and post-hoc tests, alpha was set to 0.05.

## Acknowledgements

This work was supported by NIDA grant R00 DA037271. The authors have no conflicts of interest to disclose. Special thanks to BR Cho, J Gerena, AN Herrera Charpentier, DI Olekanma and K You, for their invaluable comments on the manuscript.

## Notes

https://github.com/aneeshbal/SimpylCellCounter

