## Supplementary Method for "SimpylCellCounter: An Automated Solution for Quantifying Cells in Brain Tissue"

**SUPPLEMENTARY METHODS**

***Convolutional Neural Network (CNN) Description***

We formulated a novel approach to detecting overlapping cells in any given non-circular contour by utilizing a convolutional neural network (CNN) classifier (**Supplementary Fig S1**). Our CNN consisted of an 8-layer network built in a Python implementation of a Keras Sequential model^1–3^. The first layer is a convolutional input layer followed by 3 down-sampling layers and 2 up-sampling layers, inspirited by the U-net architecture^4^. Following the convolutional layers, the output is flattened and is inputted into a 32-neuron, fully-connected dense layer. Lastly, this 32-neuron layer fully connects to a 3-neuron, dense layer which makes the final classification for number of cells^5^.

The network was trained to classify either 1, 2 or 3 cells in any individual contour. The training and validation data consisted of 30,000 fabricated images of cells, each image containing either 1, 2 or 3 cells on a 100 x 100 pixel square. By doing this, the network was exposed to many examples of 2 or 3 overlapping cells and was able to appropriately classify them. The network was trained on Google’s Colab GPU services for 15 epochs (24,000 samples per epoch), achieving a training accuracy of 96.10% and a validation accuracy of 90.50%. It should be noted that a sub-95% validation accuracy is to be expected on such a task given that if two cells were perfectly overlapping, then it would be nearly impossible to classify the image accurately. Though the CNN is pre-trained and highly-optimized for SCC, the code for retraining the CNN or modifying its architecture is available at: <https://github.com/aneeshbal/SimpylCellCounter/blob/master/recreationFunctions/CNN_for_Overlap.ipynb>

***OCFU Threshold Standardization***

OCFU’s definition of threshold relates to the relative “darkness” of colonies compared to the background intensity with higher threshold values, resulting in a more stringent analysis. With IMJM and SCC, the threshold value strictly indicates the pixel intensity at which a binary threshold will be applied. Therefore, we aimed to standardize OCFU threshold values to IMJM and SCC. To achieve this we created 255, 12-pixel radius circles on a white background that each varied by one pixel intensity (0-255). Then, we iterated through OCFU threshold values starting from 0 to 255 and determined how many cells were counted. The number of counted cells at any given OCFU threshold value represented the equivalent threshold value in IMJM and SCC. Therefore, we were able to accurately compare relative threshold values across all automated methods (**Supplementary Fig S2**).

**SUPPLEMENTARY FIGURE LEGENDS**

**Supplementary Figure 1: Neural Network Design.** SCC utilizes an 8-layer convolutional neural network (CNN) to classify overlapping cells in non-circular contours. Classification begins by SCC extracting non-circular contours following filtering in step 3 (**Fig 1B**). Non-circular contours are pasted in the center of a 100 x 100 white background, and this constitutes the input layer to the CNN. The input layer is then fed forward to a convolutional layer while down-sampling is performed via a rectified linear unit activation function (ReLU) for 4 iterations. Then, for 2 iterations, up-sampling is performed via ReLU followed by flattening. A 32-neuron, fully connected dense layer (FCDL) then receives this flattened input and feeds it forward to a 3-neuron FCDL. This final 3-neuron layer comprises the output layer which performs the final classification. ReLU activation = rectified linear unit activation function; down-sampling = reducing dimensionality of an input to allow for assumptions about its features; up-sampling = recovers the lost resolution from down-sampling, FCDL = fully-connected dense layer where all neurons connect to all input neurons, and fully connect to their feedforward layer neurons; output neuron = final neuron activation that determines classified categories.

**Supplementary Figure 2: OCFU Threshold Standardization.** Standardization of OCFU’s threshold function, allows for accurate comparison to IMJM and SCC thresholds. X-axis represents the user-defined, binary pixel value threshold. Y-axis represents the corresponding threshold parameter for each respective method. For example, if the user selects 115 as a pixel value to threshold, then the OCFU threshold value to choose (orange line) is approximately 62. OCFU (orange), IMJM (gray), SCC (blue).
