## Supplementary figures and images for "SimpylCellCounter: An Automated Solution for Quantifying Cells in Brain Tissue"

### Supplemental Figure 1

#### SUPPLEMENTARY FIGURE S1

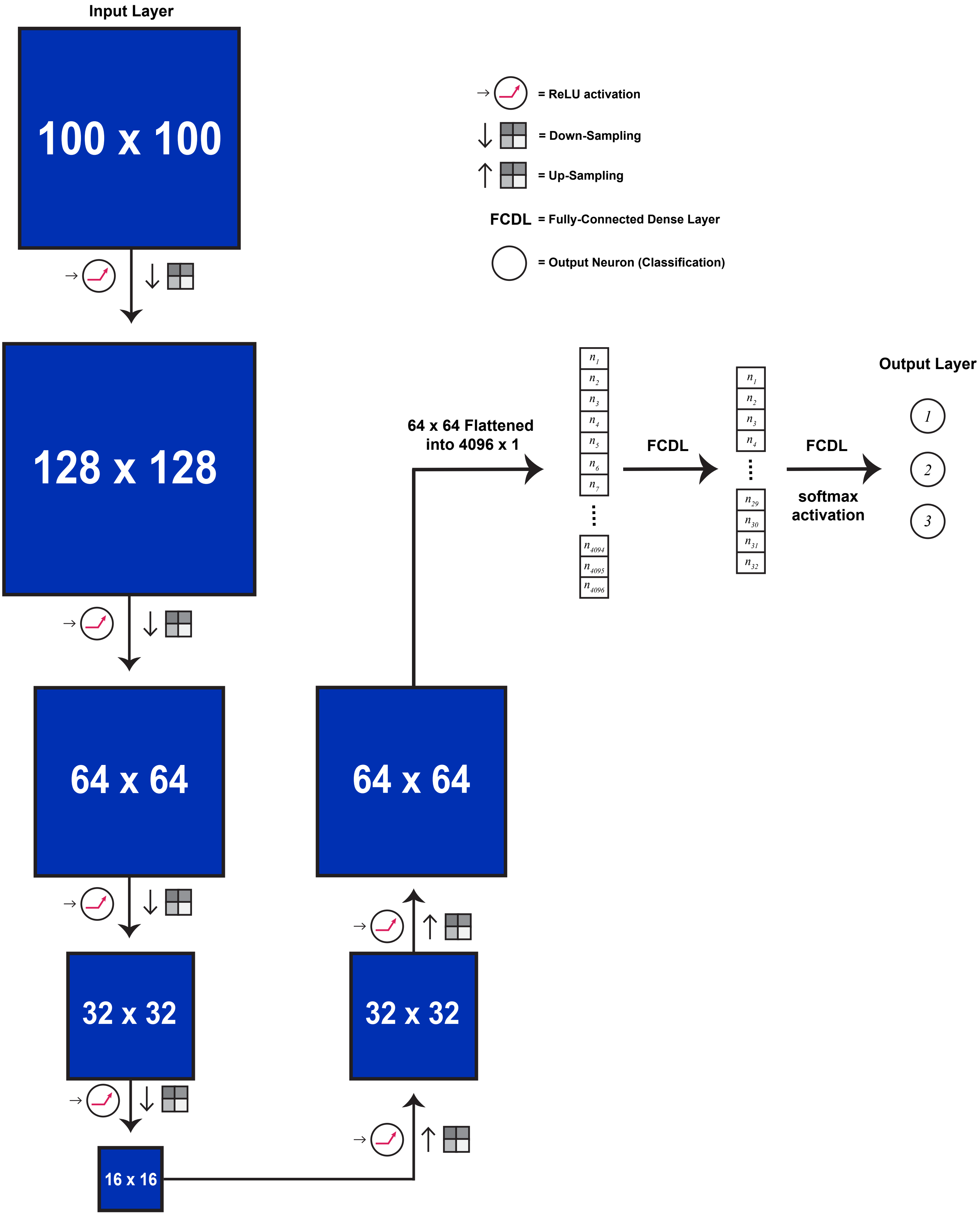

### Supplemental Figure 2

#### SUPPLEMENTARY FIGURE S2

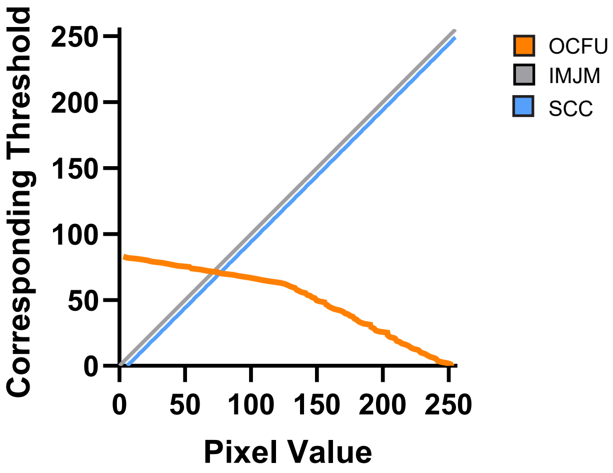
